# Effective seed sterilization methods require optimization across maize genotypes

**DOI:** 10.1101/2023.12.14.571779

**Authors:** J. Jacob Parnell, Gaurav Pal, Ayesha Awan, Simina Vintila, Gabriella Houdinet, Christine V. Hawkes, Peter J. Balint-Kurti, Maggie R. Wagner, Manuel Kleiner

## Abstract

Studies of plant-microbe interactions using synthetic microbial communities (SynComs) often require the removal of seed-associated microbes by seed sterilization prior to inoculation to provide gnotobiotic growth conditions. A diversity of seed sterilization protocols have been developed in the past and have been used on different plant species with various amounts of validation. From these studies it has become clear that each plant species requires its own optimized sterilization protocol. It has, however, so far not been tested if the same protocol works equally well for different varieties and seed sources of one plant species. We evaluated six seed sterilization protocols on two different varieties (Sugar Bun & B73) of maize. All unsterilized maize seeds showed fungal growth upon germination on filter paper, highlighting the need for a sterilization protocol. A short sterilization protocol with hypochlorite and ethanol was sufficient to prevent fungal growth on Sugar Bun germinants, however a longer protocol with heat treatment and germination in fungicide was needed to obtain clean B73 germinants. This difference may have arisen from the effect of either genotype or seed source. We then tested the protocol that performed best for B73 on three additional maize genotypes from four sources. Seed germination rates and fungal contamination levels varied widely by genotype and geographic source of seeds. Our study shows that consideration of both variety and seed source is important when optimizing sterilization protocols and highlights the importance of including seed source information in plant-microbe interaction studies that use sterilized seeds.

## Introduction

Plant-associated microorganisms are critical for plant development, evolution, and resistance to biotic and abiotic stresses (Cordovez et al., 2019; Philippot et al., 2013; Turner et al., 2013). Past studies on plant-microbe interactions have focused on studies in natural soils, which provide realistic growth conditions for the plant, but at the same time make it difficult to understand specific interactions between plant and microbes due to the high diversity of microbial communities and the presence of many biotic and abiotic factors that can confound results. In recent years there has been growing interest in studying plant-microbe interactions using plants inoculated with fully-defined synthetic microbial communities (SynComs) grown under controlled gnotobiotic conditions. These SynCom-based studies have led to major discoveries on the dynamics of plant-microbe and microbe-microbe interactions, plant colonization and community assembly processes, and the molecular mechanism leading to plant phenotype changes induced by specific microbes (Vorholt et al., 2017, Carlstrom et al., 2019; Finkel et al., 2020; Wagner et al., 2021; Wippel et al., 2021). Ultimately, the hope is that development of plant specific SynCom inoculants can improve yields and crop protection in sustainable agriculture (Shayanthan et al., 2022).

SynCom experiments often rely on sterile or gnotobiotic seeds to be able to relate the effects of specific microorganisms or groups of microorganisms to plant and microbial community responses. Seeds usually carry diverse microorganisms both on their surface as epiphytes (Torres-Cortes, 2018), and inside the seed as endophytes (Samreen et al., 2021). For gnotobiotic SynCom studies, this natural seed microbiome needs to be removed. Furthermore, removal of seed endophytes is likely necessary for experimental investigation of their importance for plant-microbiome interactions; for example, testing the hypothesis that seed endophytes have outsized effects on seedlings (Newcombe et al. 2023, 2018).

Many seed sterilization protocols have been used to generate microbe-free seeds with varying success. These include chlorine gas (Leukel and Nelson, 2940), plasma treatment (Khamsen et al., 2016), gamma irradiation (Cuero et al., 1986), heat treatment (Watts et al., 1993; Bishop et al., 1997; Regalado González, et a., 2020) microwaving (Seaman and Wallen 1967), and sodium hypochlorite/bleach (Davoudpour et al., 2020; Lindsey et al,. 2017). To help reduce seed endophytes, Watts et al (1993) proposed a pre-soak treatment prior to heat treatment. Some methods for eliminating seed contamination can negatively affect seed germination (Davoudpour et al., 2020). No consensus has emerged yet for the best sterilization protocol that balances eliminating microbes while maximizing germination. In a comprehensive analysis of seed microbiota from 50 plant species, Simonin et al. (2022) found high variation in seed community composition from the same species, possibly suggesting a variety or location effect. Most sterilization protocols are optimized for a single variety of the respective plant species and it is currently unclear if protocols for sterilization need to be adapted for different plant lineages and/or sources of seeds.

One important question when evaluating seed sterilization approaches is, how to assess the sterility of the resulting seeds. In theory, sterility implies the complete absence of viable microbes, however, currently no feasible approach exists to determine complete absence of viable microbes in sterilized seeds. Past studies have assessed sterility of seeds by directly placing seeds in or on a growth medium (Seamen and Wallan, 1967; Dawar et al. 2007; Cuero et al. 1986; Davoudpour et al. 2020; Scott et al. 2018; Watts et al. 1993; Lindsey et al. 2017), or by spreading the last wash solution on nutrient agar (Pal et al. 2022). Few studies (see Davoudpour et al., 2020) have gone beyond traditional plate cultivation to determine the presence of viable microbes following sterilization using fluorescence microscopy. While some approaches do not depend on cultivation (e.g., Bergmann et al., 2022), most do. Because many bacteria and fungi are difficult or impossible to culture, their elimination is therefore challenging to test.

Here we evaluated and compared seed sterilization protocols for maize (*Zea mays* L.), which is one of the most important cereal and bioenergy crops (Wallace et al., 2014; Nuss and Tanumiharjko, 2010; Edgerton 2009; Farrell et al., 2006, Pacala and Socolow, 2004), and for which extensive genetic and molecular tools to study plant-microbe interactions are available, including a well-established SynCom (Niu et al., 2017; Salvato et al., 2022; Krumbach et al., 2021). We compared six sterilization protocols which combine variations of sodium hypochlorite/ethanol surface sterilization, prolonged heating of seeds to eliminate endophytes, pre-soaking to eliminate endophytes, and germination with fungicide. Since, based on prior studies, we knew that fungal overgrowth is one of the main indicators of contamination of maize seeds, we used fungal outgrowth after germination as the measure to assess sterility. We determined the impact of each protocol on fungal outgrowth and seed germination rates for two different varieties of maize: a commercial hybrid sweet corn (“Sugar Bun”) and the inbred maize variety B73. We then explore the impact of the most effective sterilization protocol for B73 on fungal outgrowth and germination rates in other inbred and hybrid varieties and seeds sourced from different geographical locations.

## Materials and Methods

### Experiment 1: Testing six sterilization protocols on two maize varieties

For the first experiment in this study, we tested six sterilization protocols on two varieties of maize seeds: a commercial hybrid sweet corn variety (Sugar Bun 267T lot 68707, Johnny’s Selected Seeds, Waterville, Maine, USA) and an inbred field corn line (B73; produced by hand-pollination in a field at Central crops research station in Clayton NC, 2021). Before sterilization, seeds were visually checked and discarded if cracks or damage were visible (Gu et al., 2019). Each sterilization treatment was applied to 72 seeds of each variety.

We tested two different protocols (short and long) and two modifications of each of the two protocols (pre-soaking and germination with fungicide) (Fig. 1B). For the short sterilization protocol, quality-checked maize seeds were placed in a 50 ml conical tube (15-20 seeds/tube) and the tubes were filled to 45 ml with 70% ethanol and shaken by inversion for 3 min. Ethanol was discarded and each tube was filled with a 2% sodium hypochlorite solution and shaken by inversion for 3 min. The sodium hypochlorite solution was discarded and seeds were rinsed 5 times with sterile deionized water.

**Figure 1.**
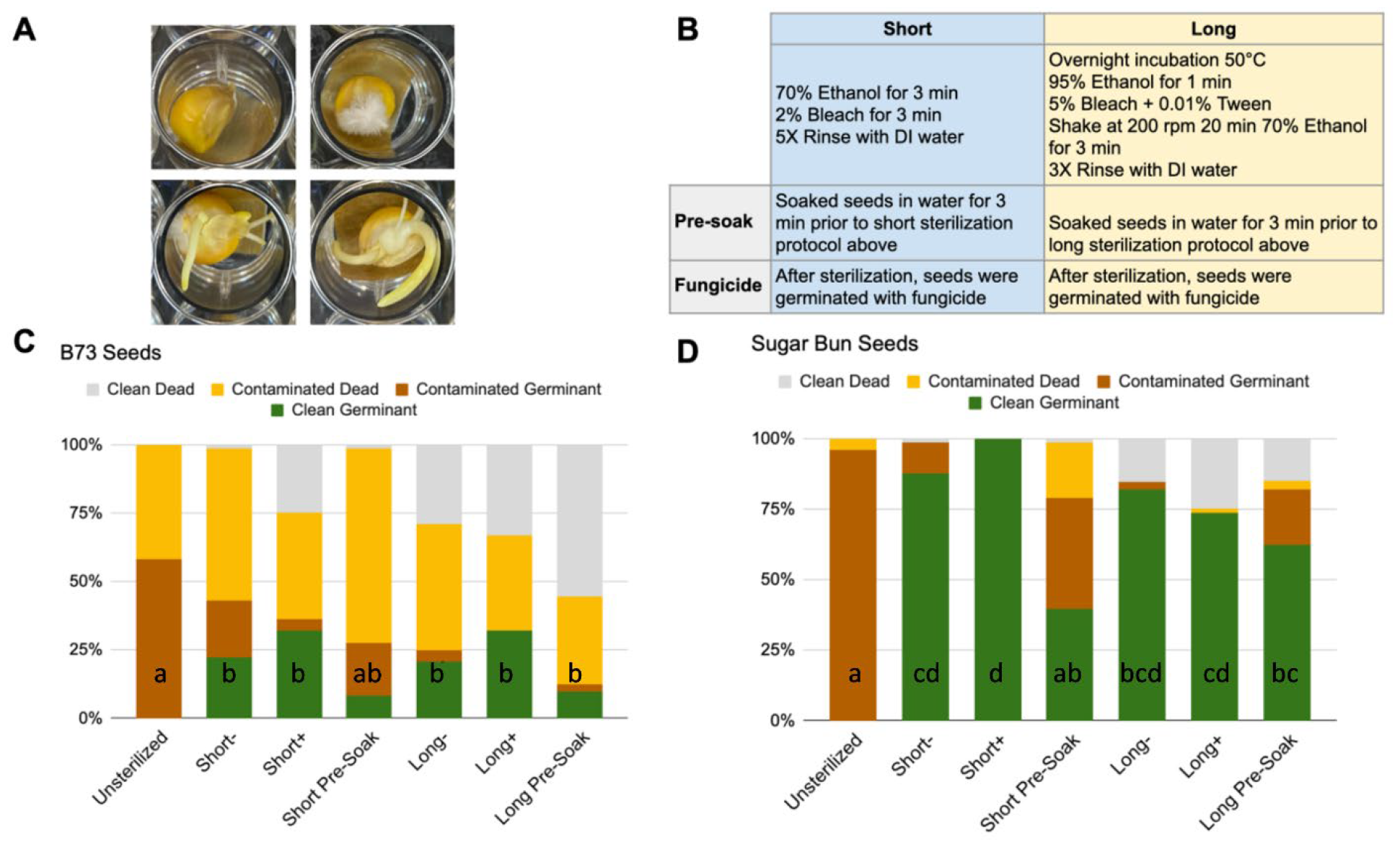
Six seed sterilization protocols tested on two varieties of maize. Examples of clean dead (top left), contaminated dead (top right), contaminated germinant (bottom left), and clean germinant (bottom right) B73 seeds (A). Overview of the six sterilization protocols used (B). Percentage of seeds that germinated and/or were contaminated with fungal growth for B73 (C), and Sugar Bun seeds (D) for each of the different protocol variations. Letters indicate significant differences for clean germinants (Tukey HSD, *p*-value < 0.05).

For the long sterilization protocol, quality-checked maize seeds were placed in aluminum foil overnight (>12h) in an incubator at 50 °C. To avoid crowding and provide even dispersal of heat, several foil packages were spread evenly in the incubator, with each package containing 10-15 seeds. After the heat treatment, 15-20 seeds were placed in a 50 ml conical tube and the seeds were immersed in 95% ethanol (∼45 ml) and shaken by inversion for 1 min. Ethanol was discarded and each tube was filled with 5% sodium hypochlorite + 0.01% Tween20 solution and placed in a shaker at 200 rpm for 20 min. The sodium hypochlorite + Tween20 solution was discarded and each tube was filled with 70% ethanol and shaken by inversion for 3 min. Ethanol was discarded and seeds were rinsed 3 times with sterile deionized water. After the third rinse, seeds were immersed in sterile deionized water for 1 h.

In addition, we examined the impact of two modifications for each of the protocols above. The first was to determine whether pre-soaking the seeds lowers the incidence of fungal endophytes as suggested previously (Watts et al., 1993). We evaluated the impact of pre-soaking by soaking the seeds for 3 min in sterilized deionized water prior to both the short (“short-pre-soak”) and the long (“long-pre-soak”) sterilization protocols.

Working next to a flame, we placed individual treated seeds into the wells of sterile 24-well polystyrene tissue culture plates (FabLab cat. No. FL7123). Wells were lined with sterile germination paper (Anchor Paper Co., USA). Three 24-well replicate plates were prepared per treatment (total of 72 seeds for each treatment) with 200 μl of sterile 0.5 X Murashige and Skoog basal salts liquid medium (Caisson Labs, Smithfield, UT, USA) per well. For the second modification, we included one treatment with the addition of a fungicide (Chlorothalonil; 1% Daconil in 200 μl of 0.5 X Murashige and Skoog) following the short (“short+”) and long (“long+”) sterilization protocols. The tissue culture plates were covered with lids, randomized, and incubated in a gravity convection incubator at 30 °C in the dark. Germination and fungal growth was recorded for each well at 7 days.

Germination was recorded as the appearance of the radicle emerging from the maize seed. Fungal contamination was recorded as any visible fungal growth. To determine the impact of contamination on germination, we recorded the proportion of seeds that did or did not germinate and were contaminated and not contaminated (Fig. 1A).

We counted each plate of 24 seeds as one experimental replicate. The six treatments for the experiment are detailed in Figure 1B. We replicated each of the six treatments three times for each of the two maize varieties (Sugar Bun and B73) and converted the raw counts of seeds germinated or contaminated to a percentage for each plate. We then normalized percentages by transforming them using a central log ratio transform (Greenacre, 2021). We tested the distribution within each treatment for normality using the Shapiro-Wilk test and for homogeneity of variances using Levene’s Test. We performed a one-way ANOVA on the normalized data to determine the relationship between the treatments and the number of uncontaminated-germinated seeds.

### Experiment 2: Testing one of the best performing protocols on four genotypes from four different sources

For the second experiment, we addressed how the best protocol for B73 in experiment 1 performed across common maize inbred and hybrid seeds sourced from different locations and years. Specifically, we applied the long sterilization protocol with germination in fungicide (long+) to B73, Mo17, and their two hybrids, B73 x Mo17, and Mo17 x B73. These two hybrids are from reciprocal crosses between identical lines-in each case the female parent is listed first. For these four genotypes we tested seed sourced from (i.e. produced by hand-pollination in) Central crops Research Station in Clayton, NC (CL) in 2021 and 2022 (CL21 and CL22), San Juan de Abajo, Nayarit, Mexico (MX; 2021-2022), and a greenhouse in North Carolina, USA (GH; 2021-2022). For the greenhouse we only had B73 and B73 x Mo17 seeds.

For the comparison of seed sources, we sterilized 24 seeds of each variety and from each source and recorded germination and contamination as described above. We used a comparison of independent proportions using a mid-P approach to Fisher’s exact test (StatsDirect, 2013) to determine differences in germination and contamination.

## Results

### Experiment 1: Best and worst performing protocols differ between two seed varieties

There was a significant effect of treatment on the number of clean germinants in both B73 (*F*(7, 16) = 5.45, *p* = 0.002) and Sugar Bun (*F*(7, 16) = 5.45, *p* = 0.002). All of the untreated (non-sterilized) seeds for both varieties showed fungal growth (Fig 1C & 1D). All treatments, except the short pre-soak, significantly increased the number of clean germinants compared to the untreated group for both seed varieties. While none of the sterilization protocols completely eliminated fungal contamination, the B73 seeds treated with the long sterilization with fungicide (long+) treatment and the Sugar Bun seeds treated with the short sterilization with fungicide (short+) treatment yielded only clean germinants (Fig.1).

Relative to the untreated seeds, the short sterilization protocol significantly increased the number of clean germinants for both B73 (from 0% to 22.2% clean germinants; Tukey HSD adjusted p-value <0.05) and Sugar Bun (from 0% to 87.5% clean germinants; Tukey HSD adjusted p-value <0.01). In both genotypes, the short sterilization protocol significantly reduced the contamination rate providing clean germinants.

Similar to the short sterilization protocol, the long sterilization protocol significantly increased the number of clean germinants for both B73 (from 0% to 20.8% clean germinants; Tukey HSD adjusted p-value <0.05) and Sugar Bun (from 0% to 73.6% clean germinants; Tukey HSD adjusted p-value <0.01). In both genotypes, the long sterilization protocol significantly reduced the contamination rate providing clean germinants.

The addition of fungicide during germination consistently increased the number of clean germinants for B73 in both the short (22.2% short sterilization to 31.9% short sterilization with fungicide) and long (20.8% long sterilization to 31.9% long sterilization with fungicide) treatments. However, these increases were not significant. For Sugar Bun seeds, fungicide addition following the short sterilization protocol increased the number of clean germinants from 87.5% to 100% (not significant). Fungicide addition following the long sterilization protocol decreased the number of clean germinants from 81.9% to 73.6% (not significant).

Pre-soaking the seeds in water prior to the short sterilization protocol did not significantly increase the number of clean germinants compared with untreated seeds in B73 seeds (8.3% clean germinants) or Sugar Bun seeds (25% clean germinants) and showed no improvement over the short or the long sterilization protocol alone.

### Experiment 2: Genotypes and seed sources differ strongly in fungal contamination and germination rates

We tested the efficiency of the long sterilization protocol with fungicide on four genotypes (B73, Mo17, B73 x Mo17 and Mo17 x B73) from four different sources (i.e. either place or production year differed), Central crops Research Station in Clayton, NC (CL) in 2021 and 2022 (CL21 and CL22), San Juan de Abajo, Nayarit, Mexico (MX; 2021-2022), and a greenhouse in North Carolina, USA (GH; 2021-2022). Seed source had a large effect on the amount of fungal contamination and germination after sterilization. The long sterilization protocol with fungicide (long+) eliminated fungal contamination in all but three of the seeds that germinated across all varieties (N = 157). Fungal contamination could not be completely removed from the NC field sourced (CL) seeds, however, most of the seeds that still showed fungal contamination after sterilization did not germinate. In contrast, none of the greenhouse (GH) and Mexico field-sourced (MX) seeds showed any contamination. The contamination rate following sterilization of CL seeds ranged from 4% to 33% and differed between genotypes (Fig. 2), however, there was no significant difference in seed contamination from year to year (CL21 and CL22) when comparing within genotypes.

**Figure 2.**
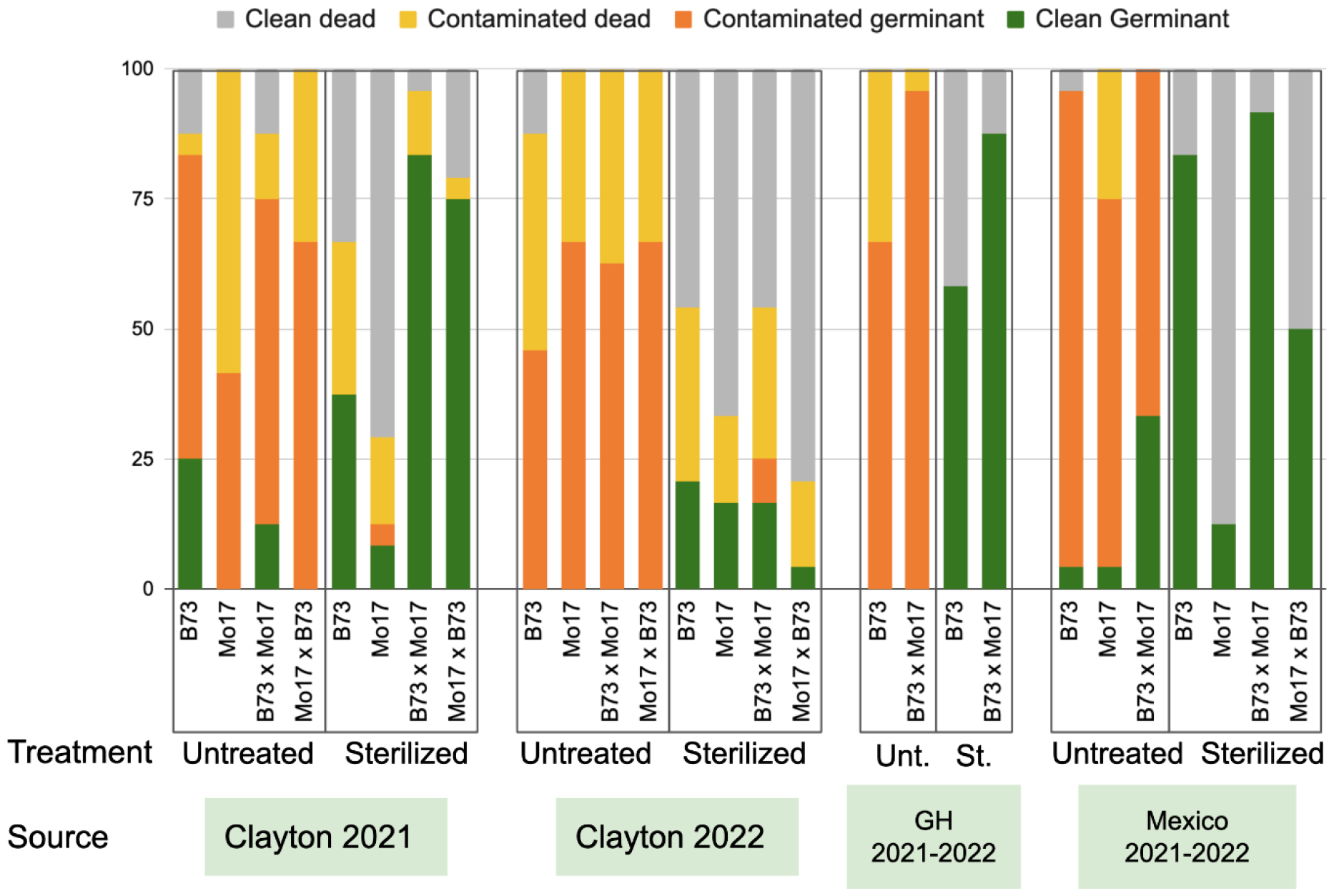
Efficiency of the long sterilization protocol with fungicide on four genotypes from four sources. Percent germination and contamination for 14 different combinations of genotype and source, sterilized and untreated. Labels are source (Clayton, GH= greenhouse, and Mexico), year or season harvested (2021 and 2022), and genotype (B73, Mo17, B73 x Mo17, and Mo17 x B73). For GH we only had B73 and B73 x Mo17 seeds. Each bar represents the results from 24 sterilized seeds.

From year to year (comparing CL21 and CL22), the inbred lines B73 (exact two sided (mid) P = 0.2259) and Mo17 (exact two sided (mid) P = 0.4272) didn’t show any difference in the proportion of germinating seeds. However, both hybrids, B73 x Mo17 (exact two sided (mid) P < 0.0001) and Mo17 x B73 (exact two sided (mid) P < 0.0001) showed significantly lower germination rates in seeds harvested in 2022 than seeds harvested in 2021. The source of seeds did not significantly impact the proportion of germinated Mo17 seeds. Mo17 seeds from all sources only had a 12.5% (SD 4.2%) average germination rate. However the source location did significantly impact the proportion of B37 seeds that germinated. The proportion of B73 seeds that germinated was not significantly different between CL21 and GH (two sided (mid) P = 0.1648), but is for CL22 and GH (exact two sided (mid) P = 0.0099). The germinating proportion of CL seeds was significantly lower than MX seeds for both CL21 (two sided (mid) P = 0.0015), and CL22 (two sided (mid) P < 0.0001) seeds. There was no significant difference in the proportion of Mo17 x B73 seeds that germinated when comparing CL and MX sources. The proportion of germinated B73 x Mo17 seeds appears to be driven more by the year differences than source differences. The germination proportion of B73 x Mo17 seeds from the CL22 source (25%) was much lower than those from the CL21 source (83%). However, germination of CL21 B73 x Mo17 seeds did not differ from germination of GH or MX seeds (two sided (mid) P > 0.99 for both comparisons). We found a weak negative relationship between fungal contamination and germination rate (Fig. 3).

**Figure 3.**
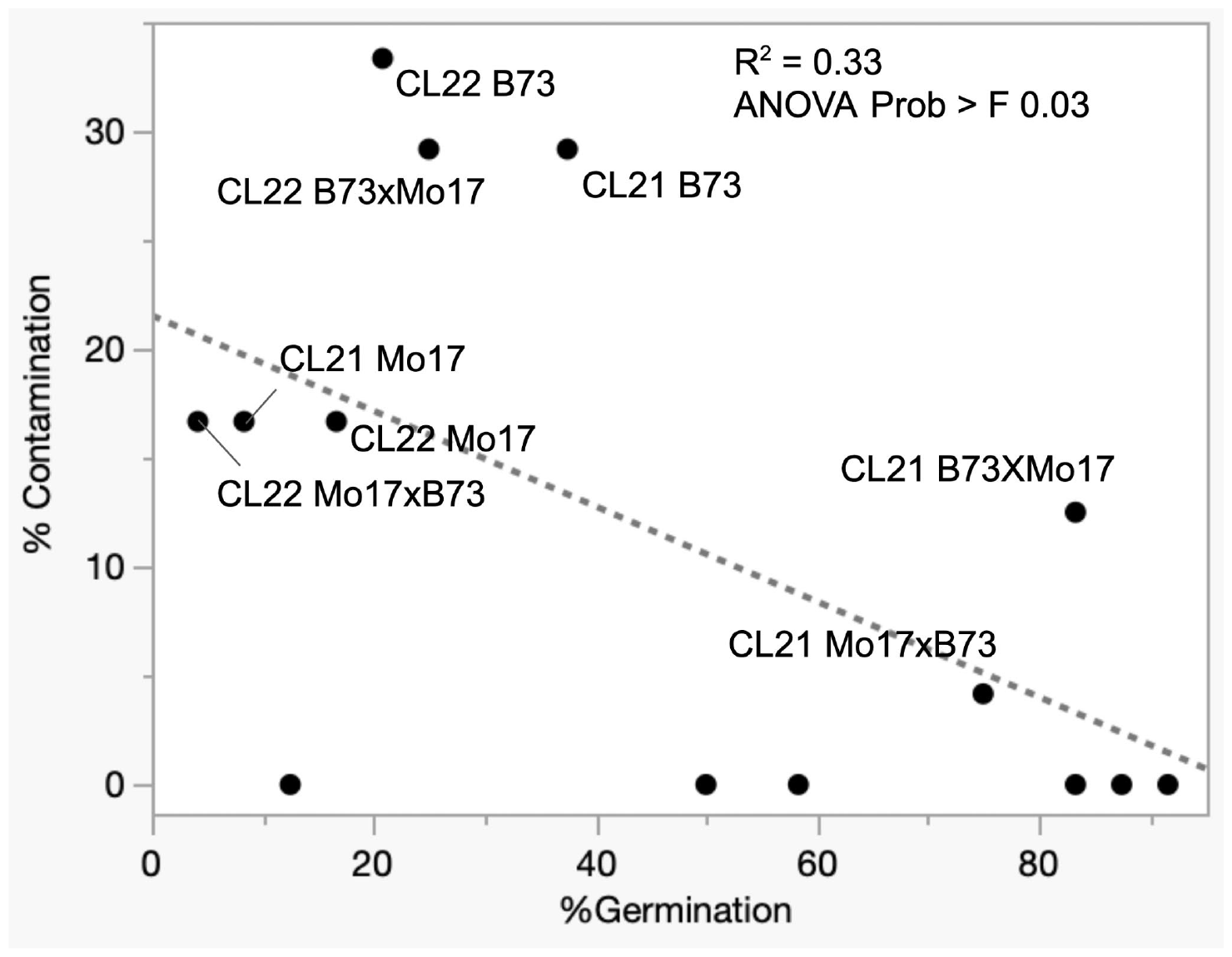
Relationship between germination and contamination. More contamination following the long+ sterilization protocol is associated with lower germination. Only Clayton-sourced seeds were included as GH and Mexico seeds show no contamination.

## Discussion

The goals of our study were (1) to identify protocols that allow for the efficient sterilization of maize seeds, while maintaining good germination rates, and (2) to test if the same protocol would work best for seeds from different varieties and sources or if the protocol would need to be adapted to the specific seeds. We found that the best performing protocol, as well as seed sterilization success differed between maize varieties and seed sources.

All of the seeds from both B73 and Sugar Bun harbored fungi, which manifested as fungal outgrowth in untreated seeds, illustrating the importance of a seed sterilization protocol when exploring plant interactions with known microorganisms in SynComs. The sterilization approaches that we tested removed up to 100% of the fungal outgrowth from Sugar Bun seeds and up to 65.3% from the B73 seeds. The short sterilization protocol with germination in a dilute fungicide yielded 100% germination with no contamination for Sugar Bun seeds. However, none of the sterilization methods tested here were able to completely remove fungal contamination from B73 seeds that were sourced from a field in North Carolina. The long sterilization protocol with fungicide did, however, eliminate all contamination from germinants.

B73 was more sensitive to sterilization protocols than Sugar Bun. The short sterilization protocol inhibited germination of B73 by almost 30% while, except for sterilization following pre-soaking, there was no impact on Sugar Bun germination. Although the long sterilization protocol lowered the germination rate of B73, compared with the untreated control, we found that using the long protocol with germination in fungicide led to a ∼32% germination rate producing clean, uncontaminated seedlings. It also appears that the long sterilization protocol mostly inhibited germination in contaminated B73 seeds that may have germinated, as there was no difference in the proportion of clean germinants between the short and long protocols with fungicide.

By extending our study to different varieties, and different source locations, we were able to show that fungal contamination levels vary greatly between seed sources even after treatment with our best-performing sterilization protocol. We found that fungal contamination was much more prevalent and difficult to eliminate in field-grown seeds from North Carolina. Seeds that were grown in the greenhouse in North Carolina or field-grown in Mexico showed no contamination following sterilization. These differences may be due to the differences in the growth environments at these locations. The CL seeds were grown in an environment with many sources of fungal contamination and no chemical control (fungicide) was used. In Mexico, a fungicide was used and in GH, we hypothesize that there were fewer sources of contamination since it was contained in a greenhouse and grown during winter. We also found that there may be some negative interaction between the fungal contamination and germination of the hybrid maize varieties used in this study as indicated by the higher contamination and lower germination of CL22 seeds versus CL21 seeds.

Our study illustrates two potential strategies for ensuring gnotobiotic growth conditions for SynCom studies in maize roots. One strategy is to pre-germinate seeds following sterilization to allow choosing of clean seedlings for inoculation and transplantation into the experimental system as has been previously done for maize (Niu et al., 2017). The second strategy, recommended when inoculation can’t wait until after germination, is to optimize a sterilization strategy for each variety, source, and year of seed used in the study to maximize reduction in fungal contamination while preserving a sufficient amount of germination.

Based on our results we can make the following recommendations for testing sterilization protocols on new varieties and sources of maize seeds. If pre-germination of seedlings is an option we would recommend to test both the short and long sterilization protocols without pre-soaking followed by germination in wells containing fungicide as at least one of these protocols will likely eliminate fungal contamination while retaining a good amount of germination. If the subsequent study will include fungal inoculants the protocols should also be tested without germination with fungicide to avoid impacts of residual fungicide on fungal inoculants. If germination without fungicide does not provide a sufficient number of clean germinants an alternative seed source/variety can be chosen, or alternatively the effect of residual fungicide on the fungal inoculant after transfer of seedlings to the gnotobiotic system can be tested to determine the size of the effect of residual fungicide if any on the inoculant.

One key limitation of our study and all previous seed sterilization studies is that testing for complete absence of live microbes from seeds is difficult if not impossible (see Introduction). We used fungal outgrowth after sterilization as the measure to evaluate success of the sterilization protocol. While we think that the elimination of fungal outgrowth likely indicates elimination of other, less visible microorganisms as well, we cannot exclude that some microorganisms remain viable in or on the seed and could grow up in the plant upon germination. For experiments where complete sterility is required, one potential route to obtain further evidence of sterility could be to characterize microbial content of larger seedlings using 16S rRNA gene amplicon sequencing for bacteria and archaea and ITS amplicon sequencing for fungi. While detection of these genes would not necessarily indicate contamination, as traces of DNA from dead cells and kit contamination (Salters et al., 2014) could lead to detection, monitoring of microbial amplicon abundances in relation to plant amplicon abundances (e.g. plastid 16S rRNA gene) across a time course of growth would provide good indication of microbial growth if the microbial signal does not get diluted during plant growth. Additionally, crushing up the seeds after sterilization followed by plating on bacterial and fungal growth media could provide additional evidence for sterility of the seeds. If bacterial growth were detected, antibiotics targeting bacteria could be used in addition to fungicide during germination as has been done previously (Pal et al., 2022).

While we only evaluated protocols on maize seeds in this study, we do think that the observed variability in germination rate and contamination level after sterilization of different varieties and seed sources, will likely also be relevant for other plant species. Therefore, testing and optimization of sterilization protocols in other plant species should include multiple varieties and seed sources as well. Additionally, our data also show that for plant-microbiome studies reporting of the seed source (location and year it was harvested) is critical in addition to information on the variety to be able to assess if observed differences between studies/experiments are potentially due to differences in microbial content of seeds.

## Author contributions

All authors contributed to the conception and design of this study. JJP, GP, PBK, SV and GH ran experiments. JJP and AA analyzed the data. JJP and MK wrote the first draft of the manuscript. All authors contributed to manuscript revision, read, and approved the submitted manuscript.

## Acknowledgements

We would like to thank Mary May for technical support.

